# Codon Usage Bias Levels Predict Taxonomic Identity and Genetic Composition

**DOI:** 10.1101/2020.10.26.356295

**Authors:** Bohdan B. Khomtchouk

## Abstract

In this study, we investigate how an organism’s codon usage bias levels can serve as a predictor and classifier of various genomic and evolutionary features across the three kingdoms of life (archaea, bacteria, eukarya). We perform secondary analysis of existing genetic datasets to build several artificial intelligence (AI) and machine learning models trained on over 13,000 organisms that show it is possible to accurately predict an organism’s DNA type (nuclear, mitochondrial, chloroplast) and taxonomic identity simply using its genetic code (64 codon usage frequencies). By leveraging advanced AI and machine learning methods to accurately identify evolutionary origins and genetic composition from codon usage patterns, our study suggests that the genetic code can be utilized to train accurate machine learning classifiers of taxonomic and phylogenetic features. Our dataset and analyses are made publicly available on Github and the UCI Machine Learning Repository (https://archive.ics.uci.edu/ml/datasets/Codon+usage) to facilitate open-source reproducibility and community engagement.

## 1 Introduction

The coding DNA of a genome describes the proteins of the organism in terms of 64 different codons that map to 21 different amino acids and a stop signal. Different organisms differ not only in the amino acid sequences of their proteins, but also in the extents in which they use the synonymous codons for different amino acids. The inherent redundancy of the genetic code allows the same amino acid to be specified by one to five different codons so that there are, in principle, many different nucleic acids to describe the primary structure of a given protein. Coding DNA sequences thus can carry information beyond that needed for encoding amino acid sequence. Thus, one may ask: is it possible to classify some properties of nucleic acids from the usages of different synonymous codons rather than, with much greater computational effort, from individual nucleotide sequences themselves?

Here, we describe an attempt to classify codon usage in terms of viral, phageal, bacterial, archaeal, and eukaryotic lineage, as well as classifying codon usage by cellular compartment (nuclear, mitochondrial, and chloroplast DNA in their respective organelles). We find that several machine-learning classifiers allow one to make such distinctions reliably from genome-wide codon usage alone.

By performing secondary analysis of existing genetic datasets stored in the Codon Usage Tabulated from Genbank (CUTG) database [6], we demonstrate that genomic and evolutionary features can be learned using machine learning methods and used for identifying kingdoms and DNA types of new genomes. However, these machine learning algorithms, such as k-Nearest Neighbors (k-NN) [1], Random Forests (RF) [2], Extreme Gradient Boosting (XGBoost) [3], Artificial Neural Networks (ANN) [4], and Naive Bayes (NB) [5], each demonstrate unique model performances with different degrees of efficiency and accuracy. Therefore, we set out to compare and benchmark these algorithms in predicting the genetic composition and taxonomic identity of organisms from different taxa.

## 2 Data

### 2.1 Data Description

We examined codon usage frequencies in the genomic coding DNA of a large sample of diverse organisms from different taxa tabulated in the CUTG database. Specifically, we compile the individual files of the CUTG database (labelled ‘qbxxxspsum.txt’, xxx = vir, phg, bct, pln, inv, vrt, mam, rod, pri) into a joint database of 13028 genomes that we make available in the UCI Machine Learning Repository: https://archive.ics.uci.edu/ml/datasets/Codon+usage. For the purposes of the analysis presented in this paper, we then performed the following additional operations on this UCI dataset:

1. Discard genome entries comprising less than 1000 codons (from the ‘Ncodons‘ column). Note that there are 69 columns in the dataset.
2. Manually curate and re-classify the genome entries of the qbbct.spsum.txt file as either ‘arc’ (archaea), ‘plm’ (bacterial plasmid), or ‘bct’ (eubacteria), guided by the first word of each CUTG species name (the genus in most cases).
3. Re-classify and harmonize genome entries from the files ‘qbxxx.spsum.txt’ (where ‘xxx’ is one of ‘pln’, ‘inv’, ‘vrt’, ‘mam’, ‘rod’, or ‘pri’) as ‘euk’ (eukaryotes).
4. Identify the DNA type of the eukaryotic genomes as either 0 (nuclear), 1 (mitochondrion), 2 (chloroplast), 3 (cyanelle), 4 (plastid), 5 (nucleomorph), 6 (secondary endosymbiont), 7 (chromoplast), 8 (leukoplast), 9 (NA), 10 (proplastid), 11 (apicoplast), 12 (kinetoplast). Remove any rows that are not 0, 1, or 2 (in other words, avoid any DNA types specified by the integers greater than 2).
5. Transform CUTG codon numbers into codon frequencies by dividing them by the total number of codons of the genome entry. Note that this has already been done to the CUTG dataset that we posted on the UCI ML Repository.
6. Exclude the genome entries classified as ‘plm’ (mostly to avoid imbalanced classes in our machine learning models described in next section, since there are only 18 plasmids).

The resultant dataset then consists of 12964 organisms of which 126, 2918, 6868, 220, and 2832 belong to the archaea, bacteria, eukaryote, bacteriophage, and virus kingdoms, respectively. When categorized by DNA type, the dataset includes 9249 ‘nuclear’, 2899 ‘mitochondrial’, and 816 ‘chloroplast’ entries. The file is organized in a header line followed by one line for each genome entry (separated by ‘newline’). Items in a line are separated by one ‘comma’.

The header line provides the column headers ‘Kingdom’, ‘DNAtype’, ‘SpeciesID’, ‘Ncodons’, ‘Species Name’, followed by the three-letter specifiers of the 64 different codons (e.g., ‘AUG’), in the same order as presented in the CUTG files.

The ‘Kingdom’ column classifies the genome as either ‘vrl’, ‘phg’, ‘arc’, ‘plm’, ‘bct’, ‘pln’, ‘inv’, ‘vrt’, ‘mam’, ‘rod’, or ‘pri’, following the ‘xxx’ specifier in the CUTG file names.

The ‘DNAtype’ column contains an integer in the range 0-12 (as described above). The ‘SpeciesID’ column is the integer that denotes the species of the genome in the CUTG file. The ‘Ncodons’ column gives the total codon count in the genome entry. The ‘Species Name’ column gives the descriptive species name as in the CUTG files. Codon frequencies are given as decimal fractions (5 digits).

To facilitate code re-use/reproducibility in the greater AI/ML/data science community, we open-sourced R programming scripts detailing our machine learning analyses on Github: https://github.com/Bohdan-Khomtchouk/codon-usage

## 3 Methods

In this study, our goal is to examine whether machine learning classifiers provide optimal solutions for predicting the taxonomic identity and genetic composition of a species, which we achieve through two major steps: 1) Computation of the 64 codon usage frequencies of each species as the feature variables representing the organism; and 2) Performance comparisons between five machine learning classifiers including the k-Nearest Neighbors (k-NN), Random forests (RF), XGBoost (XGB), Artificial Neural Networks (ANN), and Naive Bayes (NB) algorithms.

Our approach also divides the classification tasks by the kingdom class and DNA types. In other words, two model performance results are shown respectively using the codon frequencies, 1) to classify the kingdom class of the organisms, and 2) to classify the three DNA types including genomic, mitochondrial, and chloroplast. For each of these classification tasks, we only include observations from categories for which there are more than 100 observations per category. As a result, the dataset includes a designated five-category kingdom class and the three-category DNA-type class.

For model performance measures, the dataset is split by 80% training and 20% testing sets. We built five machine learning classifiers for each of these classification tasks, and the model performance results are presented using five-fold cross-validation and confusion matrices on the training data. Using the caret package in R (v3.5.2)[7], we also compared the statistical metrics including the overall accuracy, precision, recall, micro and macro-F1 scores, and the AUC (area under the ROC curve) to evaluate the efficiency and accuracy of each classifier.

### 3.1 Statistical Scores

In order to effectively evaluate our proposed machine learning models, we utilized the following quantitative metrics to measure the performance of each classification task.

The *accuracy* measures the percentages of sample objects that are correctly classified and labeled. It denotes the ratio of the total number of true predictions (true positives and true negatives) to the sum of all observations. *TP*, *FP*, *TN*, *FN* each represents true positives (observations that were correctly classified as positives), false positives (observations that were incorrectly classified as positives), true negatives (observations that were correctly classified as negatives), and false negatives (observations that were incorrectly classified as negatives).

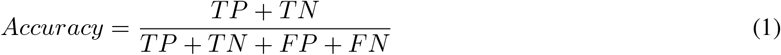

The *precision* measures the amount of variance and uncertainties of the data not explained by the fitted values of the model. The precision ranges from 0 to 1. There is often a trade-off relationship between the Precision and the Recall.

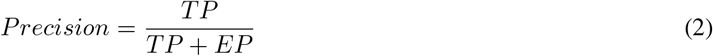

The *recall*, also known as the Sensitivity or True Positive Rate (TPR), suggests the proportion of true positives relative to the sum of true positives and false negatives (also known as the actual positives). The recall value ranges from 0 to 1, and this fraction indicates the percentages of samples or observations that are correctly classified.

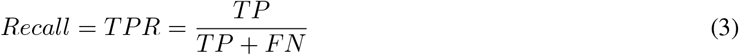

While both *precision* and *recall* offer us insights about the classification model’s behavior, their harmonic mean (F1 score) becomes a more popular metric for both binary and multiple classification situations. The *F1 score*, also known as the F-measure, is the harmonic mean of precision and recall (sensitivity)[9]. Numerically, the F1 score ranges from 0 to 1. *F*1 = 1 indicates perfect classification, which is equivalent to no misclassified samples *FN* = *FP* = 0, as shown in Equation 5. The *micro-F1 score* and *macro-F1 score* are micro-averaging and macro-averaging procedures.

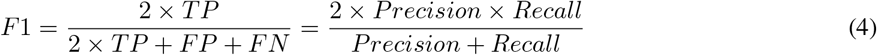

As variants of the F1 score, the *micro-F1 score* (*F*1_*micro*_) and *macro-F1 score* (*F*1_*macro*_) are obtained by first calculating a Micro-and Macro-averaged Precision (*P*_*micro*_ and *P*_*macro*_), as well as the Micro- and Macro-averaged Recall (*R*_*micro*_ and *R*_*macro*_). Here, we need to calculate the confusion matrix for every class, in which *C* = 1, 2*, ..., i, ..., n* denotes the total (n) number of classes.

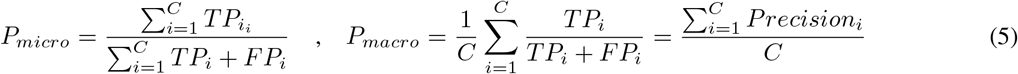

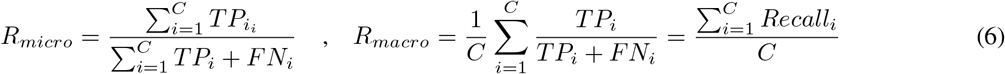

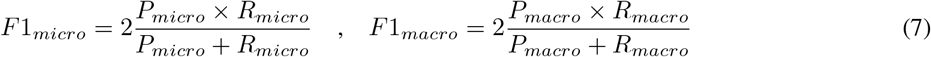

The *AUC* represents the hypothesized area under the ROC Curve. It reflects how well the model distinguishes between classes and provides a measure of performance of the ROC curve [8]. The ROC curve plots the True Positive Rate (TPR) against the False Positive Rate (FPR). The higher the proportion to 1 (the curve is closer to 1), the better the fit of the ROC curve. By summing up all the rectangular areas under the ROC curve, we can use the Trapezoidal Rule as shown in the equation below, to estimate the value of AUC. Similarly, the higher AUC value to 1 represents the selected model having more capability to distinguish between correct and incorrect classes for the samples. The weaker the AUC value (closer to 0), the less capable the model is in differentiating the real class from other false classes, which relies largely on chance.

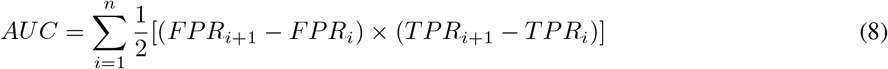

### 3.2 k-Nearest Neighbors

The k-Nearest Neighbors (k-NN) is a non-parametric classification algorithm based on classifying similar objects that cluster together in an *n*-dimensional feature space. In this paper, we used the default method of the Euclidean distance, a distance (or dissimilarity) metric to compute the pairwise differences between data observations. The Euclidean distance is the most common measure of a straight-line distance between two samples. This Euclidean distance metric formula is specified below.

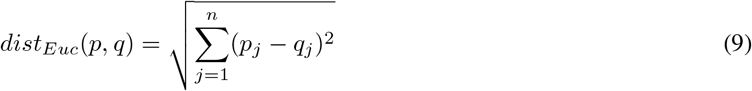

Any two objects *p* and *q* in the training set are embedded in an *n*-dimensional space (64-dimensional in our data) and the class of an object in the test set is determined to be the most common class of its closest *k* neighbors. When two objects are compared, each having *n* features, each object (*j*) is assigned to the class of neighbors (*c*) depending on the largest probability:

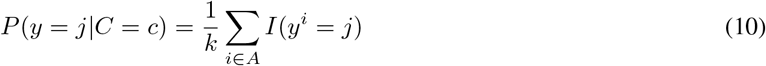

We assessed the accuracy of the model with k-th fold cross-validation procedures using a subset of our data as the training set. In the 5-fold cross-validation test, the dataset was first randomly arranged and split into 5 groups. For each unique group, this group was taken out as a test data set, and the remaining groups were assigned as the training data set. The model was developed using the four groups of the training data set, and its performance was evaluated on the test data set. This method returns a list of five accuracy values for each iteration, and the average was calculated as a cross-validation score.

We also performed the five-fold cross-validation procedure by varying the value of *k* between 1 and 5. From our analysis using different *k* options, the highest overall performance is *k* = 1, indicating that the 1-nearest neighbor algorithm demonstrates the great capability for kingdom classification tasks compared to other *k* neighbor options. The overall accuracy of the 1-nearest neighbor result is high, 0.9660, with a relatively high AUC value (0.9792) and macro-F1 score (0.9293). In the DNA-type classification task, we take the same cross-validation procedures and obtained an optimal value of *k* = 3. The overall accuracy of the 3-nearest neighbor algorithm is 0.9942, with a high AUC value (0.9997) and a good macro-F1 score (0.9867). These results indicate that the kingdoms and DNA types are distinctly clustered with very little overlaps, and the predictive models for both the kingdom classification and DNA types are robust to the training dataset.

### 3.3 Random Forests

The Random Forest (RF) classifier is an ensemble method that combines the results of different decision trees by voting across them. RF classifiers are advantageous in preventing over-fitting and handling large datasets with high dimensionality. In this paper, each decision tree arrives at a different prediction based on the predictors and the training data used, both of which are randomly chosen.

The number of trees is initialized to be 500 for building the random forest model, and arrived at reduced branches based on the random feature selection in the tree splitting process. Using 5-fold cross-validation, we tuned the *mtry* (number of variables used for splitting) parameter between 1 and 64 and arrived at an optimal value of *mtry* = 35 for the kingdom classification task with an overall accuracy 0.9298, 0.9954 in the AUC value, and 0.8611 of the macro-F1 score. Since these values are close to 1, the splitting parameter *mtry* = 35 represents a good fit for the random forest model. From the DNA-type classification results, the same steps were carried through and we obtain *mtry* = 10. A fewer number of variables are used in the DNA type classification, compared to the RF kingdom classification, yet we observe a high overall accuracy (0.9915), AUC (0.9993), and macro-F1 score (0.9832).

### 3.4 Extreme Gradient Boosting

Extreme Gradient Boosting (XGBoost) is also an ensemble method. It is a gradient boosting-based algorithm that fits an additive (ensemble) model in a forward step-wise manner with an appropriate learning rate *η*. Unlike the random forest method which builds trees independently, the XGBoost algorithm uses weak learners or shallow trees sequentially to adjust the model step-by-step. In order to optimize and minimize the loss function *L*(*y*_*i*_, *γ*) (also known as the cost function), we applied stochastic gradient descent to adjust and update our step-wise model to compute a weighted average of all sequential step-wise models *F*(*x*_*i*_). The mathematical procedures for adjusting the model sequentially are shown below. *F*_0_(*x*_*i*_) is the initial model, and *L*(*y*_*i*_, *γ*) denotes the loss function.

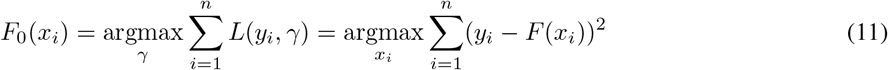

Pseudo-residuals are computed by the above equation. By running *m* number of iterations and fitting a base tree *h*_*m*_(*x*_*i*_) to these pseudo-residuals, we are able to train the model *F*_*m*_ using the training set 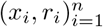.

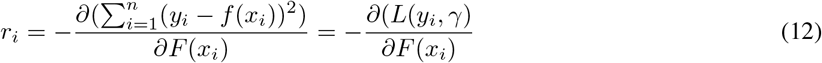

The multiplier *γ*_*m*_ is calculated as follows:

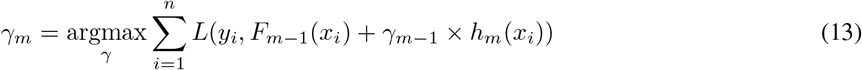

Then, we update our XGBoost model using the multiplier *γ*_*>m*_:

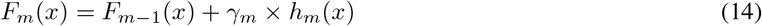

A learning rate of *η* = 0.01 is initialized and we vary the subsample ratio of the columns from 0.5 to 0.7. The maximum depth of the trees is adjusted accordingly (*max_depth* = [1, 10]). From the kingdom classification results, the best model demonstrates a high accuracy of 0.9502, a high AUC (0.997) and a macro-F1 score (0.8846). Similarly, the DNA-type classification results yielded 0.9938 in the overall model accuracy, 0.9997 of the AUC value, and 0.986 in the macro-F1 score.

### 3.5 Artificial Neural Networks

The Artificial Neural Network (ANN) algorithm consists of layers of interconnected neurons. It is also an ensemble method which learns complex relations between the codon frequency inputs, and uses layered structure to pass down inputs to the outcome. In the ANN classifier, the weights of these inter-connections are taken into account in order to determine the class of a species target. Similar to the RF classifier, ANN can powerfully handle high-dimensional datasets with large variable inputs, but also capture the shape and complex relations between the input layers.

In the kingdom classification analysis, a feedforward ANN is trained with 9 neurons in the hidden layer that yields an accuracy of 0.9132, an AUC value of 0.9901, and a macro-F1 score of 0.8425. Similarly, the DNA-type classification analysis also utilized a model with 9 neurons in the hidden layer, which provides high overall accuracy (0.9915), a high AUC (0.9997), and a solid macro-F1 score (0.9813).

### 3.6 Naive Bayes Model

Unlike other classifiers, the Naive Bayes Model is a probabilistic classifier based on the Bayes’ Theorem. It is extremely fast in model training relative to other classification algorithms. This is because the Naive Bayes Model has no complicated iterative parameter estimations, which leads to high efficiency and practical use even for high-dimensional data classification tasks. In the equation below, *P* (*class|x*) denotes the posterior probability, which determines which class the sample data belongs to. The numerator *P* (*class|x*) represents the probability that a sample would belong to a given class, and *P* (*class*) is the prior probability of a given class. The denominator at the bottom is the sample’s prior probability.

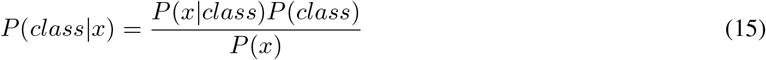

Regarding kingdom classifications, the Naive Bayes model yields an overall accuracy of 0.7522, and 0.7032 in the macro-F1 score. On the other hand, the model for the DNA type classification provides a relatively higher accuracy (0.9390) and macro-F1 score (0.8910).

However, an important assumption of the Naive Bayes model is that all the predictor variables are independent across the sample data. This assumption is not valid in this case, since we cannot assume complete independence across the codon frequencies for biological reasons. The AUC value for the ROC curves in the kingdom and DNA type classifications are 0.7980 and 0.9400 respectively, which are much weaker and indicative of worse model fit than the four aforementioned classifiers. Here, we perform the Naive Bayes classifier as a ‘worst-scenario’ case for classifier comparisons, since the Naive Bayes model relies on an assumption that is difficult to meet and often resulted in biased posterior probabilities in the analysis. Therefore, the Naive Bayes classifier is the least optimal solution for predicting the taxonomic identity and genetic composition of the organismal species in this study.

## 4 Results

A comparison of the four selected classifiers is presented below. The Naive Bayes Classifier is omitted, due to the violation of the independence assumption, which could result in potentially biased posterior probability calculations. Hence, the comparisons between the four aforementioned classifiers (k-NN, RF, XGB, and ANN) are critical when determining the outcome of the study.

The area under the ROC (Receiver Operating Characteristic) curve is measured as the AUC value below each diagram. These ROC curves describe a trade-off relationship between the TPR (true positive rate) and FPR (false positive rate) for the chosen classifier. A classifier that presents ROC curves closer to the top left corner denotes a good fit and high AUC value (closer to 1).

In Figure 1, the ROC curves are compared between k-NN, RF, XGB, and ANN classifiers for kingdom classification analysis. Both the Random Forest and XGBoost classifier show smooth ROC curves closer to 1, uniformly across the five kingdom classes. The ROC curves in the 1-Nearest Neighbor and ANN classifier give a smaller y-axis value (TPR) indicating lower true positives and higher false negatives. As in Table 1, although the precision and macro F1-scores are highest in the k-NN model, the AUC values for RF and XGB are higher (0.9954 and 0.9970, respectively), denoting a better fit in the ROC curves.

**Figure 1:**
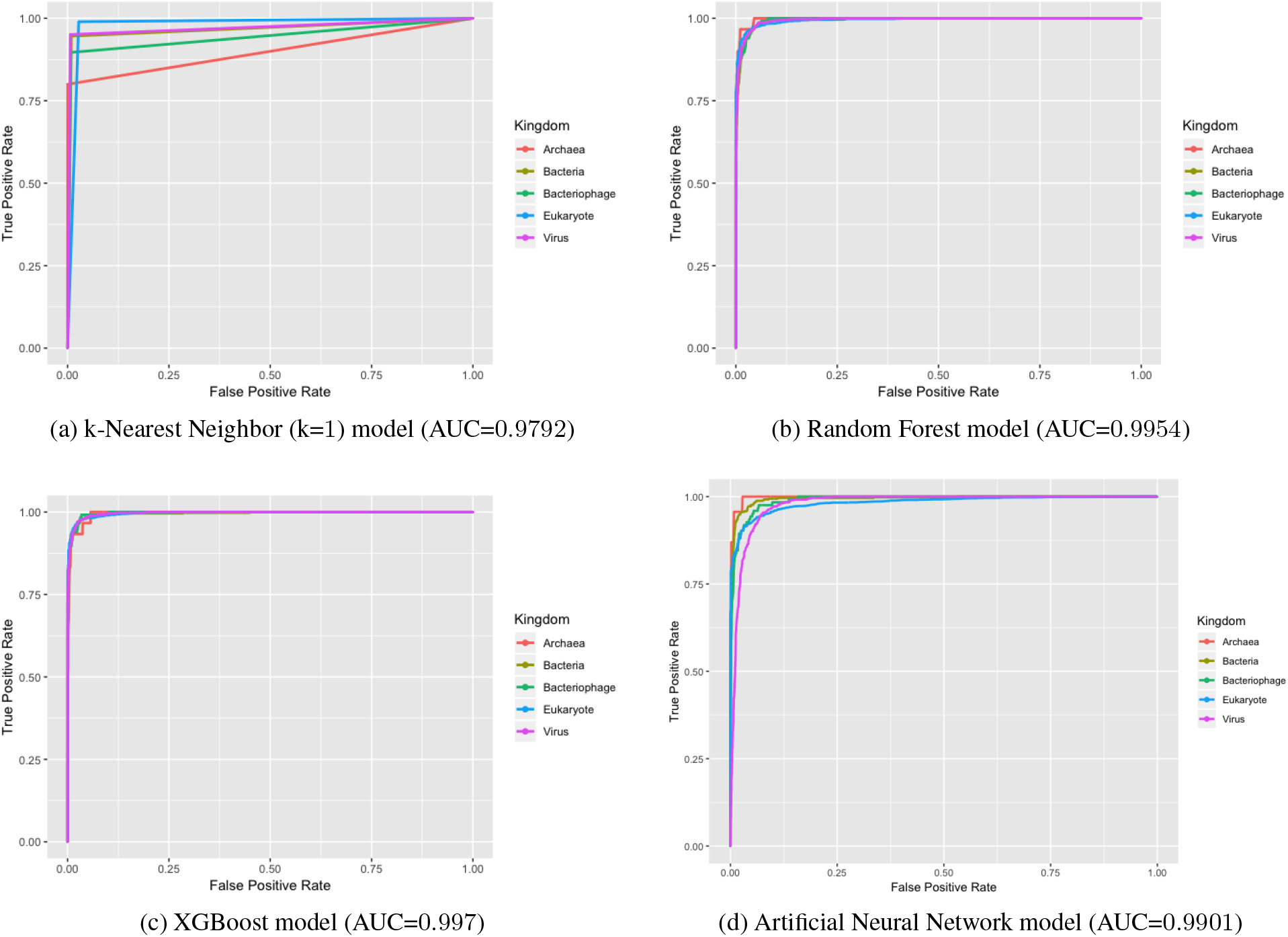
AUC ROC curves for the Kingdom Classification task

**Table 1:** Kingdom Classification Results

| Model | Precision | Recall | Micro F1-Score | Macro F1-Score | Accuracy | AUC |
| --- | --- | --- | --- | --- | --- | --- |
| k-Nearest Neighbors | 0.9660 | 1 | 0.9827 | 0.9293 | 0.9660 | 0.9792 |
| Random Forests | 0.9298 | 1 | 0.9636 | 0.8611 | 0.9298 | 0.9954 |
| Extreme Gradient Boosting | 0.9502 | 1 | 0.9745 | 0.8846 | 0.9502 | 0.9970 |
| Artificial Neural Networks | 0.9132 | 1 | 0.9546 | 0.8425 | 0.9132 | 0.9901 |
| Naive Bayes | 0.7200 | 0.3529 | 0.4737 | 0.5487 | 0.6561 | 0.8410 |

In Figure 2, the ROC curves for the four classifiers (k-NN, RF, XGB, and ANN classifiers) are compared for the DNA type classification analysis. All four ROC curve diagrams are similarly close to the top left corner, denoting a close-to-1 AUC value for these four classifiers. This indicates a good model fit and performance among the four machine learning classifiers in predicting the DNA types of the species. Table 2 gives detailed performance metrics of the four classifiers. We found that all four classifiers yield high precision and overall accuracy (> 0.99) values. In particular, the k-NN (3-NN in this case), RF, and XGB classifiers indicate exceptionally high micro and macro F1-Scores. Hence, in the DNA classification analysis, these three classifiers are of optimal choice for predicting the genetic composition of a selected organism according to its codon usage frequencies

**Figure 2:**
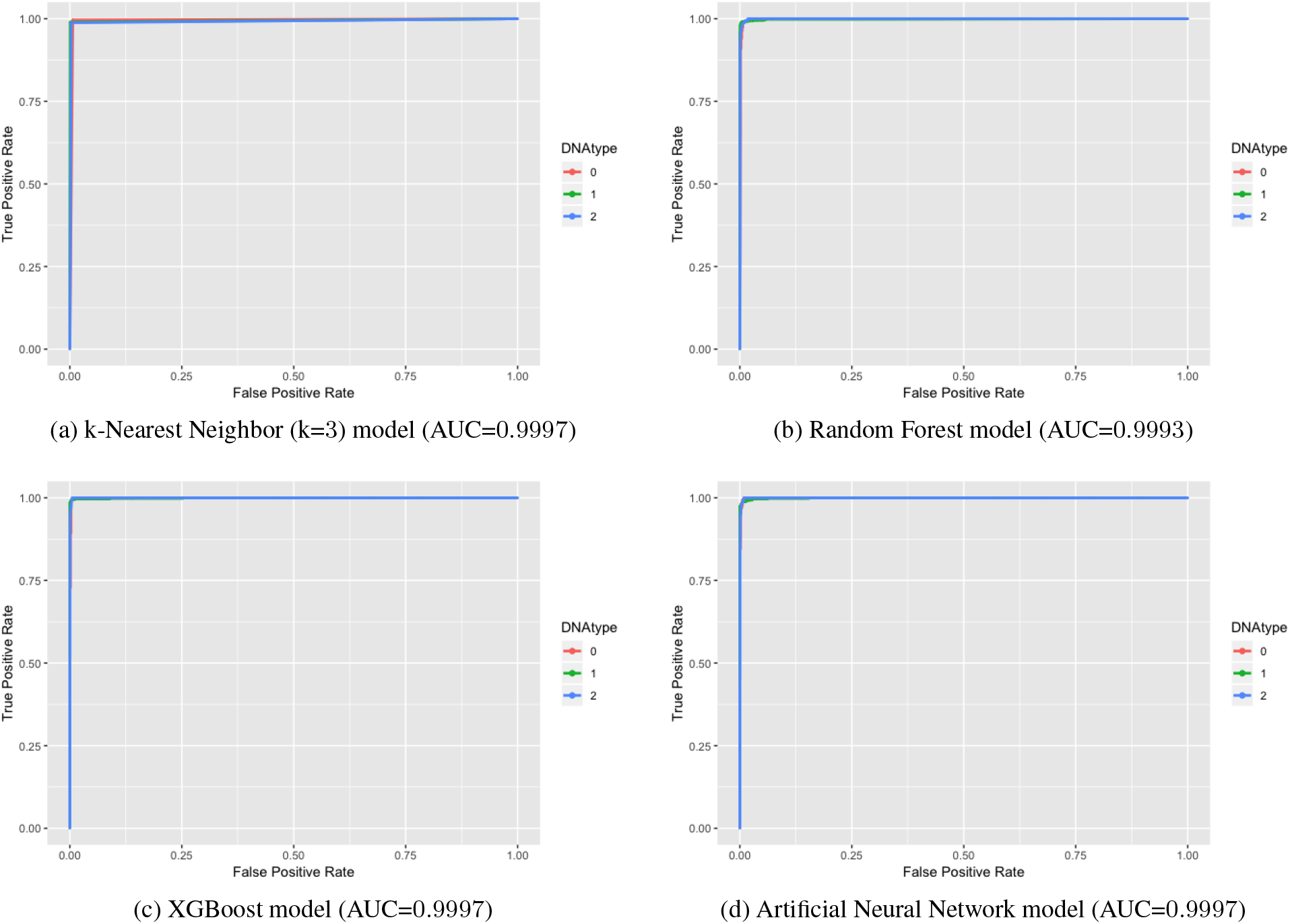
AUC ROC curves for the DNA type Classification task

**Table 2:** DNA type Classification Results

| Model | Precision | Recall | Micro F1-Score | Macro F1-Score | Accuracy | AUC |
| --- | --- | --- | --- | --- | --- | --- |
| k-Nearest Neighbors | 0.9942 | 1 | 0.9971 | 0.9867 | 0.9942 | 0.9997 |
| Random Forests | 0.9915 | 1 | 0.9957 | 0.9832 | 0.9915 | 0.9993 |
| Extreme Gradient Boosting | 0.9938 | 1 | 0.9969 | 0.9860 | 0.9938 | 0.9997 |
| Artificial Neural Networks | 0.9915 | 1 | 0.9546 | 0.9813 | 0.9915 | 0.9997 |
| Naive Bayes | 0.9085 | 0.8897 | 0.9379 | 0.8353 | 0.8870 | 0.9400 |

Figures 3 and 4 both display heatmap visualizations of the species’ codon usage frequencies by DNA types and kingdom classes. Visualizations were made with a new prototype version of HeatmapGenerator [10] available on Github branches: https://github.com/Bohdan-Khomtchouk/HeatmapGenerator. In Figure 3, the horizontal axis denotes the codon usage frequencies of a species, whereas the vertical axis shows the kingdom class these species belong to. Figure 4, on the other hand, shows codon usage frequencies of species along with their DNA types. Tree-like structures above and beside the heatmaps are clustering dendrograms performed by a hierarchical cluster analysis using the complete linkage method featured in Shinyheatmap [11]. The intensity of the colors varied across the heatmaps, showing how similar one species is relative to its neighboring species, coupled with codon usage frequency information. Various dimensionality reduction methods are tested in Figure 5.

**Figure 3:**
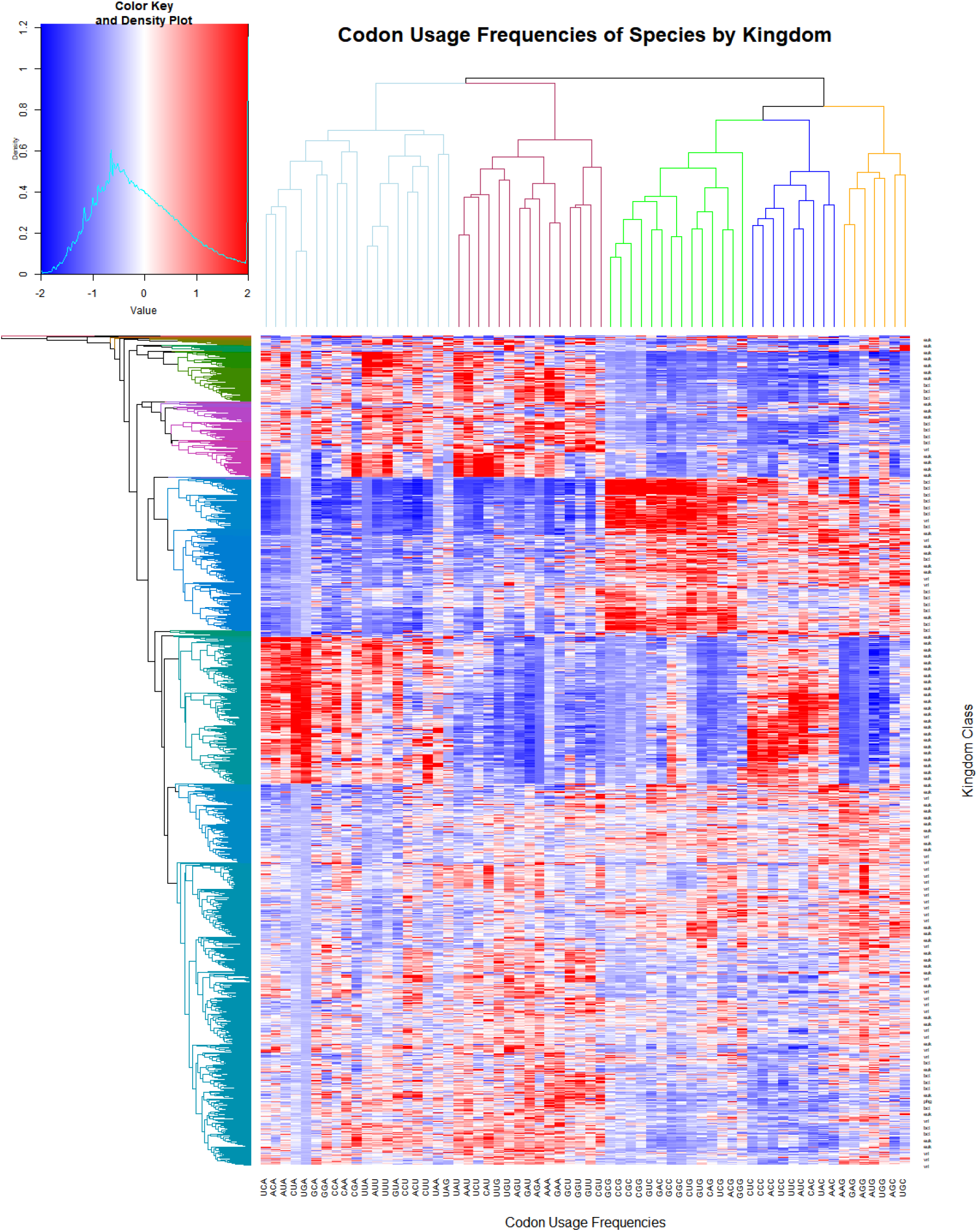
Heatmap visualization of species’ codon usage frequencies by kingdom class

**Figure 4:**
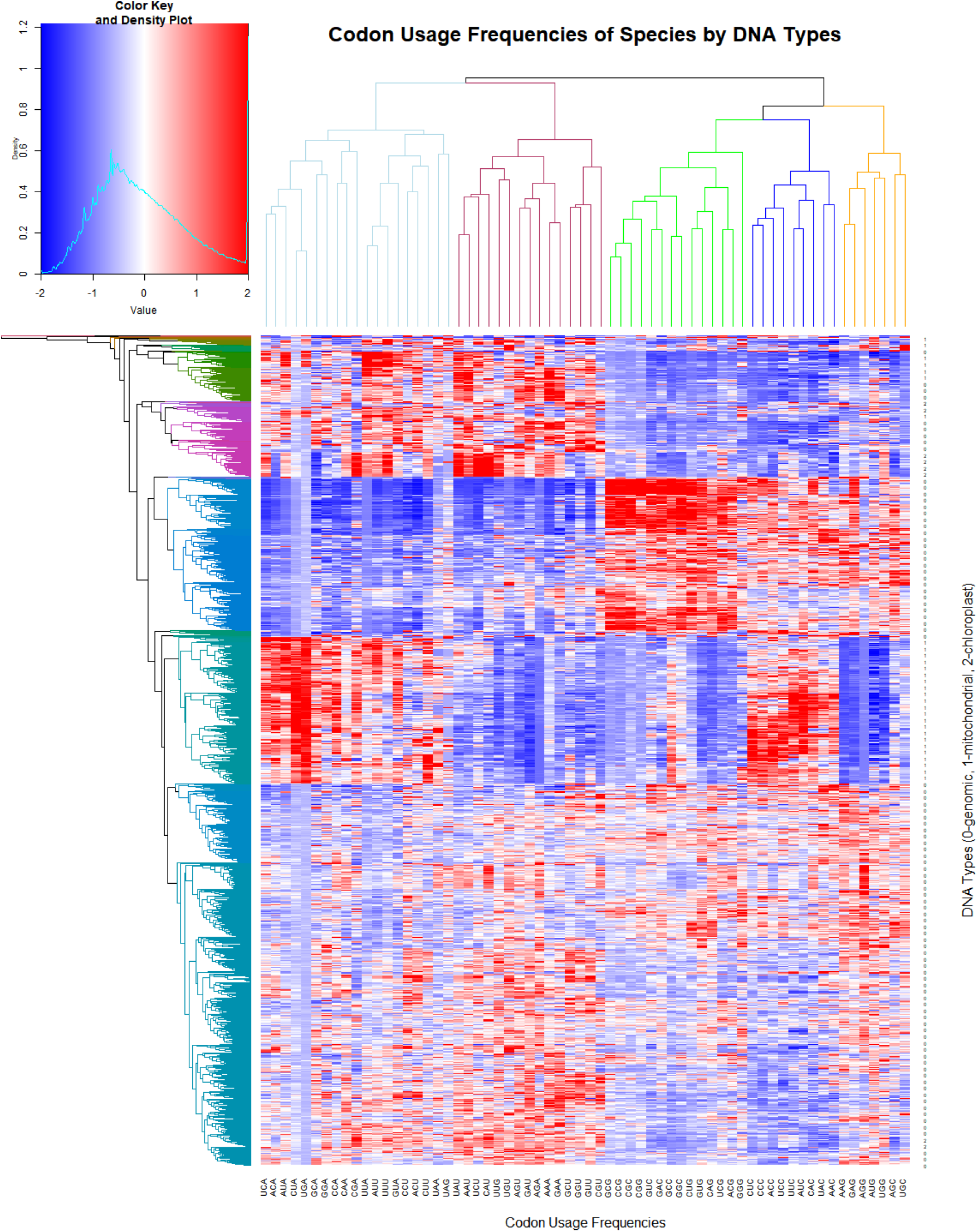
Heatmap visualization of species’ codon usage frequencies by DNA types

**Figure 5:**
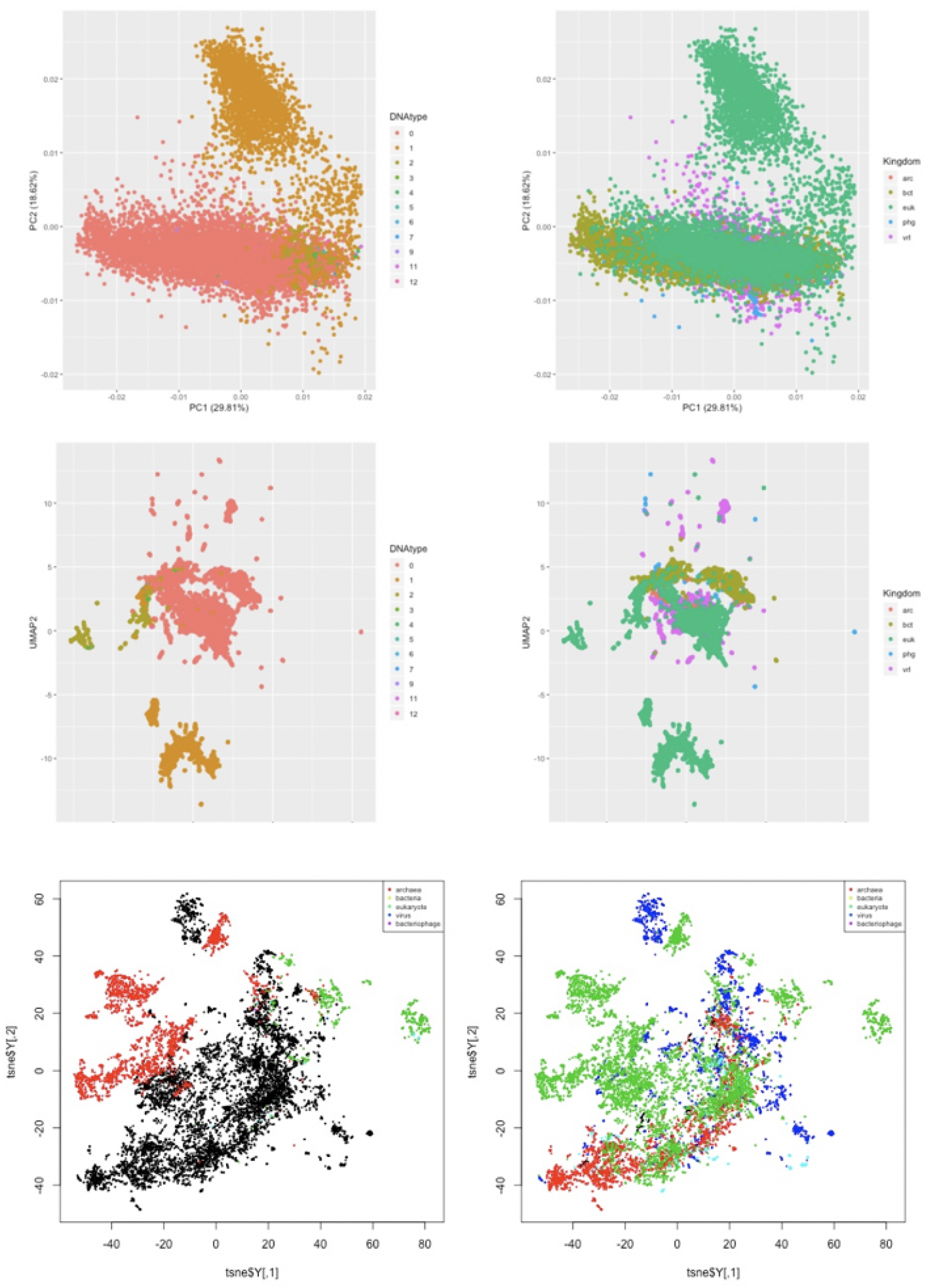
UMAP, t-SNE, and PCA plots for Kingdoms and DNAType. Not much separation observed when using Kingdoms but DNAType shows a bit of separation. Left three panels represent DNA type and right three panels represent Kingdom.

## 5 Discussion

Codon usage analyses have great potential in revealing the genetic evolution and mutation causes of living species. Future research on codon usage analyses might extend the explanation to phylogenetic lineages using a subset of characteristic codon frequencies when examining species who share similar genetic features. Furthermore, future work might apply a combined machine learning strategy, such as combining Principal Component Analysis (PCA) and other machine learning classifiers to study codon usage bias levels with a reduced codon frequency dimension. For example, we previously employed PCA and other statistical techniques to the CUTG database to find that the first principal component captures the great majority of the variation among species, and that this component reflects variation in the relative usage of A+T versus G+C nucleotides [12]. Specifically, we found an unusual pattern where very deep lineages have evolved nearly distinct patterns of codon usage – one group preferring G/C ending codons and the other preferring U/A ending codons. This fundamental pattern leads to the interesting finding that the more biased genomes show the most differences in codon usage, suggesting that variation among species is caused by horizontal gene transfer. These data explorations demonstrate that there is much that can be learned from the secondary analysis of existing genetic datasets for the study and interpretation of codon usage bias, and to facilitate such future analyses we have further consolidated, curated and harmonized individual species datasets from the CUTG and shared our new resultant master dataset on the UCI Machine Learning Repository.

## 6 Conclusions

The present findings revealed that codon usage frequencies are a shortcut to classifying DNA. These codon usage frequencies are not only useful features for predicting the taxonomic identity of a biological organism, but also powerful tools when classifying the genetic composition of life forms. From the model performances presented by five machine learning classifiers, our analysis suggests that the k-Nearest Neighbor, Extreme Gradient Boosting (XGB) and Random Forests (RF) algorithms are three optimal solutions for classifying kingdom classes during the secondary analysis of existing genetic datasets. Although the k-Nearest Neighbor (k-NN) classifier presents the highest precision (0.9660) and overall accuracy (0.9660) of the model performance, the XGB and RF classifiers yield better model fit with higher AUC (0.9970 and 0.9954, respectively) in their ROC curve diagrams. Regarding the prediction of organisms’ DNA types, all four classifiers (k-NN, XGB, RF, and ANN) show high precision (> 0.99) and overall accuracy (> 0.99) of model performance, with k-NN, XGB, and RF demonstrating high micro and macro F1-scores. In particular, the ROC curves of XGB (AUC> 0.99) and RF (AUC> 0.99) consistently show optimal performance and model fit in both kingdom and DNA-type classifications. Hence, the Extreme Gradient Boosting and Random Forest algorithms can be used to build robust and effective classifiers for identifying and predicting the taxonomic and genetic features of organisms.

## 7 Funding and Acknowledgements

The author(s) received no specific funding for this work so it will remain as a preprint posted on bioRxiv until further notice to avoid open access publication charges. BBK thanks Yunzhen (Amy) Liang and Aswathy Ajith for email correspondence discussions and assistance in early stages of the project. Also, special thanks to Dr. Wolfgang Nonner for valuable suggestions and feedback on the manuscript writing and editing process as part of previously completed work [12].

